# Attraction of *Triatoma infestans* (Klug) to adhesive yeast-baited trap under laboratory conditions

**DOI:** 10.1101/674473

**Authors:** Miriam Cardozo, Federico G. Fiad, Liliana B. Crocco, David E. Gorla

## Abstract

Existing methods to detect domestic triatomines have low sensitivity. As early house infestation detection is epidemiologically important, the exploration of better methods is required. Hence, we measured the attractiveness of a yeast-baited trap to adults and nymphs of *Triatoma infestans*, under laboratory conditions.

The assays were conducted in an experimental arena, with an experimental and a control traps placed at opposite sides and one refuge in the center area. Insects where released and the number of triatomines in the yeast and control traps were counted, after 3, 6 and 24 hours of the beginning of the experiment. We use generalized linear models within a multimodel inference approach to model the number of insects in the trap, using insect age classes, time after assay initiation and date of the experiment as predictors.

Our results show that the attraction to CO_2_ depends upon the life stage of the insects. During the 24 hours of experiment a constant number of adults were attracted to the yeast trap, while nymphs show attraction only up to the first three hours after the initiation of CO_2_ liberation. Undoubtedly, the orientation response to chemical cues deserves further studies to be fully understood.

## Introduction

Chagas disease is considered one of the most important endemic diseases in Latin America, affecting approximately 5–6 million individuals. The disease is caused by *Trypanosoma cruzi (*Trypanosomatidae), which not only infects humans but also more than 100 species of domestic and sylvatic mammals and can be transmitted by over 150 species of triatomines (Triatominae, Reduviidae) (WHO 2015).

*Triatoma infestans*, characterized by its high adaptive capacity to domestic environments, is the vector with the greatest epidemiological importance in the Southern Cone countries of South America (Rabinovich 1972; Lent and Wygodzinsky 1979).

The maximum geographical expansion of *T. infestans* distribution occurred between 1970 and 1980 with an estimated occupation area of 6.28 million km^2^, including Argentina, Bolivia, Brazil, Chile, Paraguay, Peru and Uruguay. The Southern Cone Initiative, coordinated by the Pan American Health Organization to control the transmission of Chagas disease in Latin America from 1991 interrupted the vector transmission of *T. cruzi* in Chile, Uruguay and Brazil through insecticide-based vector control, health education and house improvement programs (Dias et al. 2002). The Initiative produced a significant reduction of the distribution area of *T. infestans* to less than 1 million km^2^ (Schofield et al. 2006). Nevertheless, in arid Gran Chaco areas of Argentina, Paraguay, and Bolivia, reinfestations of human dwellings continue to occur in several provinces or departments (Ceballos et al. 2011; Bustamante-Gómez et al. 2016).

Although this species has long been found almost exclusively in domiciliary and peridomestic environments, a significant increase in the number of wild population found in sylvatic environments was reported, mainly in the Inter-Andean Valleys of Bolivia, in the Gran Chaco of Argentina, Bolivia and Paraguay (Noireau et al.1997; Rolón et al. 2011) and in a Metropolitan region from Chile (Bacigalupo et al. 2010). Recent studies also evidenced the presence of gene flow between sylvatic and intra-peridomestic *T. infestans* populations in Argentina (Piccinali et al. 2011), suggesting that sylvatic populations may be involved in the reinfestation observed in some places.

The prevention of Chagas disease depends on the elimination of the domestic colonies of triatomines. Insecticide residual spraying is very effective, but re-infestation of treated dwellings is frequent. Early detection and elimination of triatomine reinfestation is critical for long-term control; however, current methods used for vector-detection have low sensitivity. A number of alternatives have been evaluated (Abad-Franch et al. 2011), either for the detection of domestic triatomine species (like *T. infestans*), or other secondary vector species that frequently invade domestic and peridomestic structures (Cavallo et al. 2016; Cecere et al. 2016; Giraldez et al. 2016). A number of methods for triatomine detection have been tested in different ecotopes (sylvatic, domestic and peridomestic environments). Current method for routine entomological surveillance used by vector control programs in Latin America is the fixed effort active search, sometimes using a flushing out agent. Although widely adopted, it depends heavily on operator experience and has low sensitivity when vector abundance is low (Gürtler et al.1999).

Passive bio-sensors such as the Gómez-Nuñez box or Maria sensors (Gómez-Nuñez 1965; Wisnivesky-Colli et al. 1987) were tested with poor results because of methodological and operational limitations and low sensitivity to detect colonization (Pinto Dias et al. 2005). The dissection of microhabitats in which triatomines feed and shelter, such as tree holes, palm crowns, bromeliads, rock piles, burrows and bird nests, has been effective to capture sylvatic specimens, but it requires important sampling efforts, human resources and sometimes generates negative impact on the environment. Light traps have the disadvantage of attracting only hungry adults, although it has been possible to capture some species that are otherwise difficult to collect (Noireau and Dujardin 2001; Vazquez-Prokopec et al. 2004). Traps with animal bait, such as the Noireau adhesive trap, which uses a mouse as bait, were successfully used in sylvatic ecotopes (Abad-Franch et al. 2000; Gürgel-Gonçalves et al. 2003; Noireau et al. 1999). However, it is expensive due to host maintenance and some authors reported that its efficiency depends upon the triatomine species studied and the biotic region of study (Reyes-Novelo et al. 2012). As none of the explored methods show reasonable sensitivity, there is a need to develop a sensitive detection method for entomological vigilance.

Two types of non-live baited traps were also evaluated based on semiochemicals (Rojas de Arias et al. 2012) and yeast. The host orientation behavior of triatomines is controlled by physical and chemical signals, including olfactory clues such as carbon dioxide. The carbon dioxide released by the *Saccharomyces cerevisiae* cultures, is a chemical signal indicative of a food source for hematophagous insects and therefore it can evoke both behavioral responses: activation and attraction of the triatomines to the source (Lazzari et al. 2013; Guerenstein and Lazzari 2009).

Several authors have demonstrated the effectiveness of yeast traps to attract and capture *T. infestans* under both laboratory (Guerenstein et al. 1995; Barrozo and Lazzari 2004, 2006) and natural conditions (Lorenzo et al. 1998, 1999; Bacigalupo et al. 2006). Other studies have also demonstrated that yeast traps are a useful tool for the detection of potential new sylvatic habitats of *T. infestans* as well as other triatomine species and they can be a suitable alternative for their control (Bacigalupo et al. 2006; Botto-Mahan et al. 2002). Previous studies using yeast traps for triatomine detection evaluated the device during a fixed time, and generally using nymph triatomines.

Within the exploration efforts to find a method that improves the detection sensitivity of triatomines, we report a study that measured the temporal variation in attractiveness of an adhesive yeast trap for adults and nymphs of *T. infestans* under laboratory conditions.

## Materials and methods

### Experimental setting

The study was designed as an experiment involving the release of insects in an arena containing a baited trap (with yeast) and a non-baited trap (without yeast, control trap).

The experimental arena measured 100 × 80 × 80 cm, with kraft paper ground and non-climbable walls. Control and yeast traps were placed at opposite sides of the arena and folded paper as an artificial shelter (15 × 10 cm) was placed in the center.

During the assay, 8 insects were released in the area center and only after the insects had hidden in the folded shelter the traps were placed in the arena. After 3, 6 and 24 hours of the beginning of the experiment, the number of triatomines captured in the yeast trap, those adhered to the control, the loose ones in the arena and those that remained hidden in the refuge were counted. Each time the number of bugs was recorded in the traps, the replacement of the attached bugs was made, making sure that there were always 8 individuals in the experimental box.

Some tested insects were used up to 2 times with a difference of at least ten days between tests to ensure independence.

We performed seventeen series of assays for adults of *T. infestans* and thirteen series of assays for 4^th^ and 5^th^ nymph instars. During the test, the triatomines were able to move freely throughout the experimental arena. Assays were conducted at room temperature (approximately 25°C ± 2°C), in darkness starting at 10.00 am and finishing 24 hours later. The position of the control and the experimental traps was changed randomly in the successive trials, to compensate for possible external asymmetries.

### Traps

The experimental adhesive yeast traps (Figure 1) consisted of a plastic container of 500 cm^3^, with a perforated cover containing a solution of 5 g of dry yeast LEVEX^®^ + 10 g of sucrose + 100 ml of water. For the purpose of this work, we used the same concentration employed by Bacigalupo et al. (2006), as this concentration proved to be effective for the capture of wild *T. infestans* colonies in field studies.

**Fig 1.**
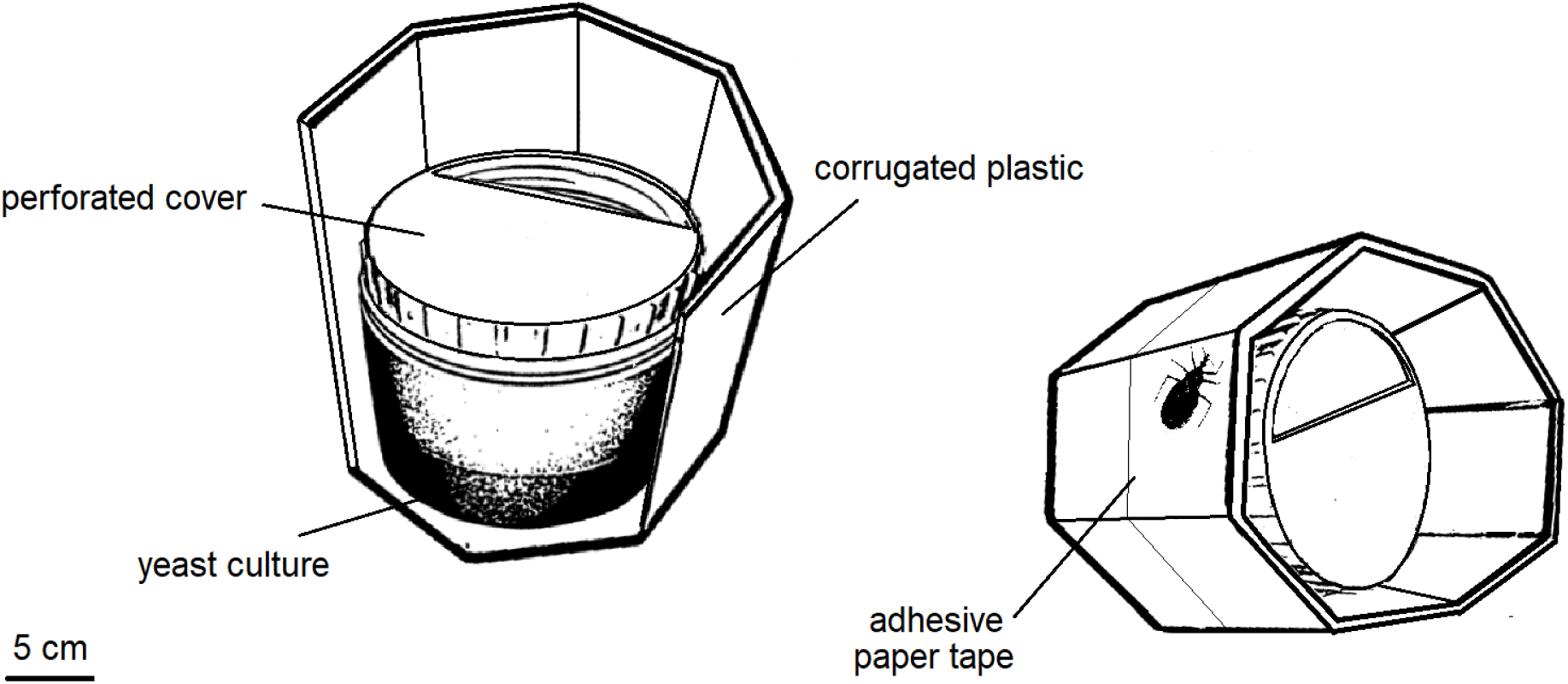

Front and side view of the yeast trap prototype used to test the attractiveness of the yeast culture for *Triatoma infestans*

The control trap had the same design of the yeast trap although it only contained a solution of sucrose (10 g of sucrose + 100 ml of water). The plastic container of both the yeast trap and the control trap was surrounded by a rectangle of corrugated plastic, covered with adhesive paper tape Doble A^®^. The corrugated plastic increases the adhesive surface and offers a refuge site. This yeast trap design is simple and cheap, easy to transport due to its low weight and volume and it can be used either horizontally or vertically so that it can be put in a great variety of sylvatic and peridomestic ecotopes.

### Insects

The triatomines used in the assays were 180 adults and 103 nymphs (83 fifth nymphal instar and 20 fourth nymphal instar) of *T. infestans*. The insects came from colonies reared during many generations (>25) in the Centro de Referencia de Vectores from the Servicio Nacional de Chagas (CeReVe-SNCh), located in Santa María de Punilla (Córdoba, Argentina).

All the triatomines were fed on chicken and kept under a natural illumination cycle under room temperature conditions (26°C ± 5°C and at 40 ± 60% relative humidity). The insects were starved for approximately 15 to 30 days before the experiments.

### Data analysis

Although the results of this type of experiment has been analyzed comparing proportions of trapped insects through ANOVA tests to detect contrast signification (e.g. Guerenstein et al. 1995), we adopted the effect estimation approach (Cumming 2012) instead of significance test. To evaluate the attractiveness of the yeast solution we used generalized linear models (GLMs) with Poisson error distribution. We consider alternative hypothesis that would explain the variation in the number of bugs captured in the yeast trap (response variable) as a function of predictor variables including the insect stages (adults or nymphs), the time intervals at which the response variable was recorded (3, 6 and 24 hours after the beginning of the assay) and the date of the experiment. The latter was considered as a temporal control to detect possible asymmetries between the assays. The candidate models considered the individual effects of each predictor on the response variable as well as joint models evaluating the additive effects and the interactions of the possible combinations. The GLMs where fitted through maximum likelihood and their relative performance were measured with Akaike’s information criterion (AICc), using the function aictab of the package AICmodavg (Mazerolle 2017) in the R software version 3.4.3 (R Core Team 2017). We assessed the significance of the effects by comparing size, unconditional standard error and 95% confidence interval. The effect-size estimates for each factor were averaged, using the modavg function from the package AICmodavg in the R software version 3.4.3 (Mazerolle 2017), for all coefficients included in models that showed a difference in AIC values ≤ 3 with the model that showed the lowest AIC.

## Results

In every replicate an average of 7.58 (95% CI [7.43;7.73]) insects, either nymph instar or adults, moved out of the central refuge. On average, 3.53 (95% CI [2.97;4.09]) adults and 2.71 (95% CI [1.97;3.46]) nymph instar that moved out of the refuge where attracted by the yeast trap, and only 5% of adults and 7% of nymph instars remained in the refuge after 24 hs.

Considering that the 8 insects released in each assay were able to choose among 4 sites (yeast trap, control trap, refuge and out of refuge), 2 is the expected number of insects to be counted at each time interval in any of these sites under the hypothesis of random selection. The median number of adults captured in the yeast trap was always > 2 (Fig. 2A), whereas in the nymph group, the median number of captured insects in the yeast trap was 2 after the first 3 hours (Fig. 2B).

**Fig 2.**
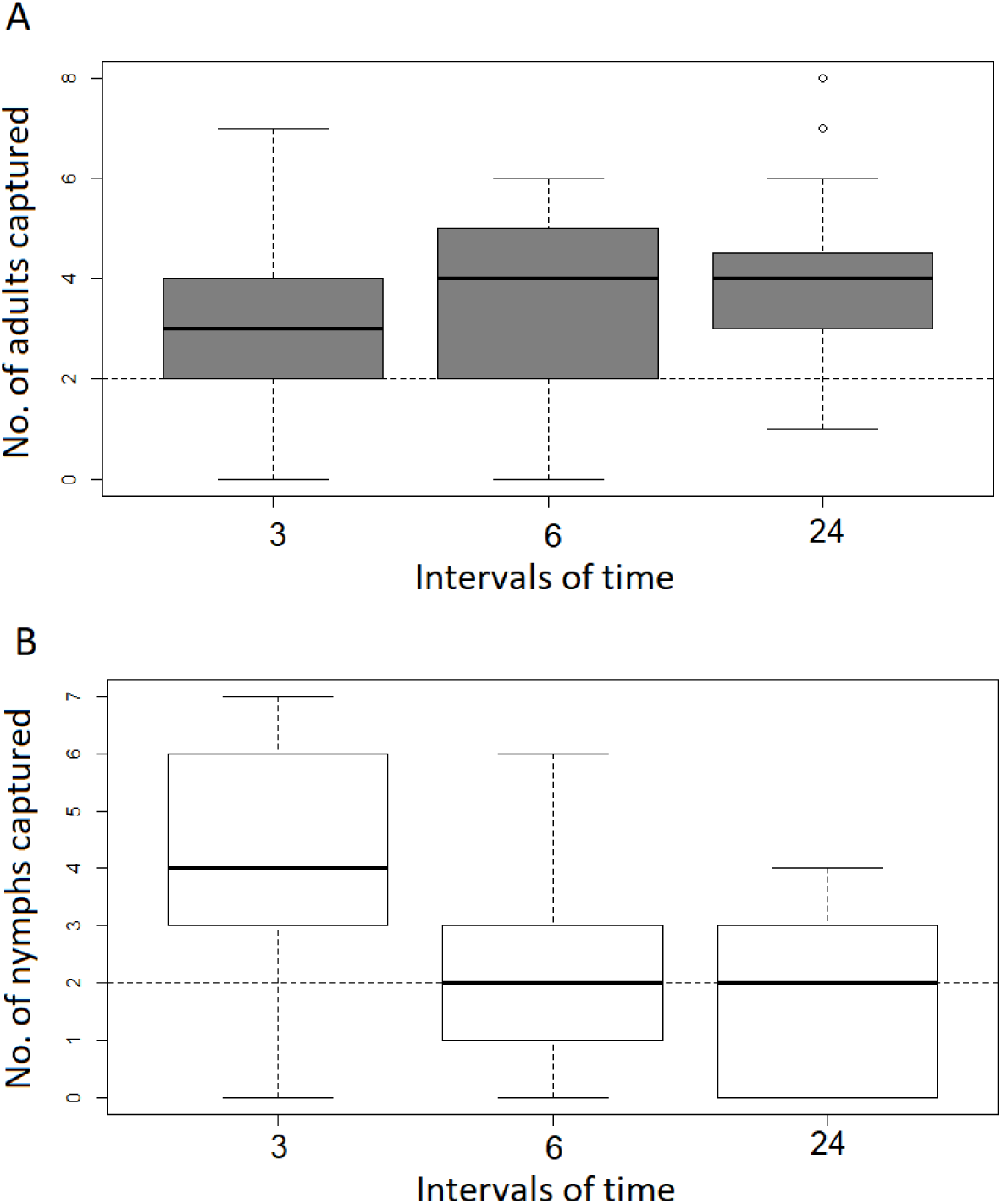

Number of *T. infestans* adults (A) and nymph instars (B) captured in the adhesive yeast trap 3, 6 and 24 hours after the beginning of the assay. The dotted line indicates the expected number of insects under the hypothesis of random selection of sites by the triatomines.

From the nine candidate models of the multimodel inference approach, three of them described equally well the results (ΔAICc ≤ 3.0) (Table 1). Two of the models included the interaction between life stage factor and the time as predictors. The presence of interaction indicates that the time effect depends on the life stage of the triatomine, meaning that attraction to the carbon dioxide source at 3, 6 and 24 hours after the beginning of the assay, differ between adults and nymphs of *T. infestans.*

**Table 1.** Model set. Exhaustive list of all GLMs considered in this study to explain variation of the number of insects recorded in the yeast trap (Y).

| Model structure | $\Delta AICc$ |
| --- | --- |
| Y (life stage + time + life stage * time) | 0.000 |
| Y (life stage + time + life stage * time + date) | 1.377 |
| Y (life stage) | 2.715 |
| Y (life stage + date) | 4.495 |
| Y (life stage + time) | 4.620 |
| Y (life stage + time + date) | 6.365 |
| Y (date) | 6.825 |
| Y (time) | 6.903 |
| Y (time + date) | 8.767 |

$\Delta AICc$ , represents the difference in the value of the Akaike's information criterion (AIC) with respect to the AIC value of the best candidate model
+, additive effect of the factor
\*, interaction effect of the factors

After detecting the life stage × time interaction, the response variable was modelled for nymph instars and adults separately using GLMs with Poisson link, considering the effects of date, time and their additive effects. Models for the nymph stage showed no effect of date (coefficient estimate = 0.01, 95%CI [0;0.01]), but a negative effect of time (coefficient estimate= −0.04, 95%CI [−0.07; −0.01]). Models for the adult stage showed no effect of date or time, and an interception estimate of 1.17 (95%CI [0.93;1.41]), meaning that number of adults remained constant during the experimental periods between 2.5 and 4.1.

## Discussion

Our results confirm the effectiveness of CO_2_ liberated by a small yeast culture to attract *T. infestans*. The detection of natural host odour blends or a single constituent (CO_2_) tends to increase the triatomine locomotor activity and trigger both behavioral responses of activation and attraction (Guerenstein and Lazzari 2009). For the first time we report the comparative attraction during 24 hours of *T. infestans* nymph instars and adults to the CO_2_ liberated by a small yeast culture. Our study shows that *T. infestans* adults and nymphs had higher locomotion activity after the experiment started than insects in similar experiments performed using other triatomine species (*T. dimidiata, T. pallidipennis T. brasiliensis, T. sordida, T. pseudomaculata*), where more than 50% of the nymphs remained in the central refuge (Pires et al. 2000; Pimenta et al. 2007), compared to 5% of adults and 7% of nymphs in the present study. The proportion of nymphs captured in the yeast trap (34%, out of the total used in the assays) was similar to that obtained for *T. infestans* in a previous study (44%, Guerenstein et al. 1995).

Our study shows that the attractiveness to the CO_2_ liberated by a small yeast culture depends upon the age classes of the insects. On average, adults were more attracted to the yeast trap than nymphs. During the 24 hours of experimental period, a constant number of adults (2.5 – 4.1) was attracted and captured in the yeast trap. However, our results showed a different behavior of nymphs, that presented attraction to CO_2_ only the first three hours of the assay, and then declining significantly over time.

Even though it has been demonstrated that the attractiveness and orientation towards CO_2_ by *T. infestans* is limited to a temporal window at the beginning of the night (Barrozo and Lazzari 2004), we observed that if a source of CO_2_ is offered, starved adults of *T. infestans* can respond to the chemical stimulus long before the beginning of the scotophase, under laboratory conditions. This behavior might be based on the possibility that the sensitivity to one specific odour is not “switched-off’ outside the temporal window associated with the search of food, as Bodin et al. (2008) suggested.

The results presented in this work add new questions about the mechanisms that modulate the CO_2_ attraction and the possible influence of vital stage of *T. infestans.*

Several studies evidence that the orientation response to stimuli is more complex than believed. This process is influenced mainly by internal factors, such as circadian clocks (Barrozo and Lazzari 2004), as well as the triatomine’s physiological state concerning its nutritional status, the proximity to oviposition and the moult cycle (Bodin et al. 2008,2009). Undoubtedly, the orientation response to stimuli and chemosensitivity deserves further studies to be fully understood. Not only to comprehend the biological basis but also because it is fundamental for the improvement of detection techniques or development of new detection tools.

The interpretation of the results of this study should take into account that the nymph’s age and the reproductive status of females were not measured. Further laboratory studies should be carried out to confirm and understand the mechanisms determining the relationship between attraction to CO_2_ and the life stage. The adhesive yeast-baited trap used in this work should be tested under field conditions to determine its sensitivity and easy-of-use before it could be recommended for the use of routine activities of triatomine surveillance.

## Acknowledgement

The authors thank Emma Bosco and Elisa Barbero for their valuable contribution with the design and development of the adhesive yeast trap. We also thank Raul Stariolo from the Centro de Referencia de Vectores from the Servicio Nacional de Chagas (CeReVe-SNCh, Santa María de Punilla Córdoba, Argentina) for the provision of the insects for experimentation. MC and DEG are supported by the National Research Council of Argentina (CONICET).

